# Inconsistencies in the published rabbit ribosomal rRNAs: a proposal for uniformity in sequence and site numbering

**DOI:** 10.1101/2024.10.11.617640

**Authors:** Swastik De, Michelle Zhou, Zuben P Brown, Raymond N. Burton-Smith, Yaser Hashem, Tatyana Pestova, Christopher U T Hellen, Joachim Frank

## Abstract

Examination of all publicly available *Oryctolagus cuniculus* (rabbit) ribosome cryo-EM structures reveals numerous confusing inconsistencies. First, there are a plethora of single nucleotide differences among the various rabbit 28S and 18S rRNA structures. Second, two nucleotides are absent from the NCBI Reference Sequence for the 18S rRNA gene. Moving forward, we propose using the Broad Institute’s rabbit whole genome shotgun sequence and numbering to reduce modeling ambiguity and improve consistency between ribosome models.

## Introduction

The translation of stored genetic information into proteins is central to all domains of life. The eukaryotic ribosome, a complex biomolecular machine composed of the small (40S) and large (60S) subunits, facilitates the reading of sequential codons in messenger RNA (mRNA) by transfer RNA (tRNA), converting this information into the language of amino acids and functional proteins. Both subunits contain multiple riboproteins bound to a backbone of different-length RNA strands: the 28S, 5.8S, and 5S rRNAs for the mammalian 60S subunit and the 18S rRNA for the 40S subunit. In recent years, while the majority of structural data for the eukaryotic ribosome has been acquired through cryo-electron microscopy (cryo-EM) and increasingly cryo-electron tomography (cryo-ET), some structures of yeast and human ribosomes have also been determined using X-ray crystallography. The data predominantly come from four species: two yeast species (*Saccharomyces cerevisiae* and *Kluyveromyces lactis*, with 262 and 14 structures, respectively), human (244 structures), and rabbit (121 structures), with all other eukaryotic organisms accounting for only 194 models (Table 1, as of 09/08/24). While the structure of the human ribosome is of high interest for understanding human biology and disease mechanisms, rabbit ribosomes are often used in their place because they can be purified in large quantities from rabbit reticulocyte lysate (RRL)—the most widely used system for in vitro analysis of translation (Jackson and Hunt, 1983) and used for reconstitution of ribosomal complexes (e.g. Hashem et al., 2013; Hilal et al., 2022). RRL is also used to prepare programmed translation reactions that are arrested at specific stages, which can then be purified for cryo-EM analysis (e.g., Bhatt et al., 2021; Shao et al., 2016; Simonetti et al., 2016, 2020).

**Table 1.** Presently available structures of eukaryotic ribosomes (as of 09/08/24, prepared using RADtool (Hassan et al., 2023)).

|  | 80S structures | 60S structures | 40S structures |
| --- | --- | --- | --- |
| **Eukarya** | 377 | 274 | 184 |
| *Saccharomyces cerevisiae* | 155 | 87 | 20 |
| *Kluyveromyces lactis* | 2 | 11 | 1 |
| *Oryctolagus cuniculus* | 64 | 21 | 36 |
| *Sus scrofa* | 11 | 7 | 3 |
| *Homo sapiens* | 80 | 89 | 75 |
| *Mus musculus* | 4 | 1 | 4 |
| *Plasmodium falciparum* | 4 | 2 | 2 |
| *Canis lupus familiaris* | 1 | 0 | 0 |
| *Triticum aestivum* | 5 | 1 | 2 |
| *Drosophila melanogaster* | 4 | 0 | 0 |
| *Spinacia oleracea* | 4 | 4 | 2 |
| *Vairimorpha necatrix* | 1 | 0 | 0 |
| *Arabidopsis thaliana* | 1 | 0 | 0 |
| *Neurospora crassa* | 4 | 2 | 1 |
| *Tetrahymena thermophila* | 1 | 4 | 6 |
| *Paranosema locustae* | 1 | 0 | 0 |
| *Thermochaetoides thermophila* | 5 | 19 | 0 |
| *Danio rerio* | 2 | 0 | 0 |
| *Xenopus laevis* | 1 | 0 | 0 |
| *Candida albicans* | 8 | 0 | 0 |
| *Spraguea lophi* | 4 | 0 | 0 |
| *Giardia spp* | 7 | 2 | 3 |
| *Encephalitozoon cuniculi* | 1 | 0 | 0 |
| *Rattus norvegicus* | 1 | 0 | 0 |

Structural investigations and extensive biochemical characterization have significantly advanced our understanding of eukaryotic ribosomal biology, providing insights into ribosome assembly (Baßler and Hurt, 2019; Broeck and Klinge, 2024), modification (Sloan et al., 2017), initiation (Hashem and Frank, 2018; Jackson et al., 2010; Querido et al., 2024), elongation (Dever et al., 2018; Djumagulov et al., 2021; Holm et al., 2023; Milicevic et al., 2024), termination (Hellen, 2018), quality control (D’Orazio and Green, 2021), and overall structure (Yusupova and Yusupov, 2014, 2017).

In this report, we investigated errors present in the rabbit ribosomal rRNA sequences within published PDB models. We also examined how these errors have affected the current structures and have prepared sequence-corrected structures for rabbit 28S, 18S, 5.8S, and 5S rRNAs.

### Discrepancies among published 28S rRNA sequences

The 60S ribosomal subunit is composed of 5S, 5.8S, and 28S rRNAs, along with various ribosomal proteins. The 28S rRNA functions as a ribozyme, catalyzing peptide bond formation essential for protein synthesis (Yusupova and Yusupov, 2014). This catalytic property is associated with RNA’s ability to fold into compact structures, creating cavities that serve as binding sites for ligands. Furthermore, studies have shown that specific elements of the 28S rRNA, such as es27l and es39l, play critical roles in binding, recruiting, coordinating, and regulating ribosome-associated protein complexes like the nascent polypeptide-associated complex (NAC) (Knorr et al., 2019; Lentzsch et al., 2024). Inconsistencies among published 28S rRNA *Oryctolagus cuniculus* sequences create issues for comparative studies and may lead to inaccurate conclusions. Aligning these sequences to a consistent reference would minimize ambiguities, improve research accuracy, and support the development of new therapies, especially for diseases like Alzheimer’s (Payão et al., 1998). Since the rabbit ribosome is a commonly used model, having accurate and widely accepted numbering is crucial for pinpointing and understanding rRNA modifications, which are significant in the context of certain cancers. For example, human ribosome modifications have been precisely mapped by multiple groups, highlighting the importance of standardized references in ribosomal research (Holvec et al., 2024; Temaj et al., 2022; Barozzi et al., 2023; Krogh et al., 2020; Cui et al., 2024). rRNA sequences are crucial for understanding ribosomal structure and function, but discrepancies between published sequences hamper research. We compared rabbit 28S Rrna sequences from various sources, including published PDB structures and template sequences from NCBI GenBank, RNA Central, ENA, and Rfam databases, to identify the discrepancies and evaluate their impact. Seven high-resolution PDB structures from different groups published as rabbit ribosomes were selected to avoid any lab-specific biases.

To identify a reliable sequence as our template, we conducted an extensive search through several sequence databases, including NCBI GenBank, NCBI RefSeq, RNA Central, Rfam, NIH Biosample, NIH Bioproject, and the European Nucleotide Archive (ENA). Initially, we suspected that the absence of a dependable 28S rRNA sequence for rabbits might have contributed to numerous sequence errors in the PDB structures. However, we identified a reliable source in GenBank entry AAGW00000000.2, which corresponds to the *Oryctolagus cuniculus* (Thorbecke inbred breed) whole genome sequencing project, released in 2005 and re-curated in August 2009. This GenBank entry is linked to multiple ENA, Rfam, and RNA Central entries, all curated as the 28S rRNA sequence of *Oryctolagus cuniculus*. The template sequence used for our study is found in RNA Central under accession code URS00009AB771_9986. This sequence originated from the Broad Institute’s rabbit whole genome sequencing trials (OryCun2.0), which employed shotgun sequencing methods to assemble the entire rabbit genome. A team of Broad Institute scientists completed a deep coverage (7x) draft of the rabbit genome, offering significantly higher accuracy compared to previous sequences. We also considered the NCBI Genome assembly UM_NZW_1.0, which is a New Zealand White rabbit sequence from the Shanghai Institutes for Biological Sciences (Bai et al., 2021). Although the New Zealand White sequence offers higher quality with 40x redundant coverage compared to the 7x coverage of OryCun2.0, we opted for the OryCun2.0 sequence for several reasons: 1) it is more thoroughly curated (providing the complete length of over 5000 nucleotides for 28S sequence compared to the ∼3600 nucleotides of the UM_NZW_1.0 sequence), and 2) it is more commonly used by the biochemical and structural groups. The availability of this high-quality, thoroughly curated 28S rRNA sequence from OryCun2.0 provided a reliable template for our comparative analysis, which was essential for accurately identifying and understanding discrepancies observed in rabbit 28S rRNA sequences reported in various PDB structures.

The comparison of published rabbit 28S rRNA sequences reveals many erroneous variations. Using the Broad Institute’s rabbit sequence as a template for alignment, we compared the seven selected PDB entries with 28S rRNAs with the template’s sequence. These comparisons revealed numerous discrepancies, ranging from single nucleotide changes to substantial insertions and deletions, indicating a lack of consistency and accuracy in reported structures. In this report, we have documented single nucleotide differences to highlight the issues arising from the lack of a validated consensus sequence. While major insertions and deletions could be tabulated, we deliberately chose not to focus on these variations due to various complications associated with them. Instead, we emphasized on the need for a standardized consensus sequence, which is crucial for accurate structural data interpretation across rRNA variants. Without such a sequence, alignment discrepancies are inevitable, and each variant would have a unique and inconsistent numbering system beyond the first indel. We chose to omit a detailed discussion of major indels to maintain clarity and focus on the necessity of a unified reference framework.

The number of single nucleotide changes varies from none to as many as forty-seven (PDB 6P5I) (Acosta-Reyes et al., 2019). The extent of these variabilities suggests that the inconsistencies are caused by a combination of factors, such as using different methodologies, starting models, and sequence correction practices.

The origins of the starting models play a crucial role in determining the final sequence accuracy. For instance, PDB 6GZ5 from the Spahn group matches the template rabbit sequence, showing no nucleotide changes. This high level of sequence accuracy may be attributed to a careful sequence correction process conducted by the authors, who thoughtfully edited the starting human ribosome model (PDB 5AJO) to align with the rabbit sequence (Behrmann et al., 2015). This approach underscores the importance of thorough sequence verification and correction, especially when working with models from closely related species. In contrast, the 6P5I PDB entry from our own lab (Acosta-Reyes et al., 2019) exhibited several single nucleotide mutations, deletions, and insertions, likely due to the initial use of a *K. lactis* ribosome structure (PDB 5IT9/5IT7) as a starting point (Murray et al., 2016). This example highlights the potential challenges that can arise when using models from more distantly related species without comprehensive sequence validation.

Interestingly, many of the PDB entries we analyzed trace their origins back to a common progenitor PDB entry, 3JAH, a structure of a rabbit ribosome published by the Ramakrishnan lab (Brown et al., 2015). This entry was itself derived from another rabbit ribosome model, PDB 3J92 (Shao et al., 2015), which in turn was based on an earlier boar ribosome model, PDB 3J7O (Voorhees et al., 2014). Ultimately, these sequences can be traced back to a single human ribosome model, 4V6X, published by the Beckmann group (Anger et al., 2013) (Fig. 1). This lineage illustrates how a single starting model can influence subsequent studies, and it suggests that any inaccuracies or assumptions in earlier models may persist through multiple iterations.

**Figure 1.**
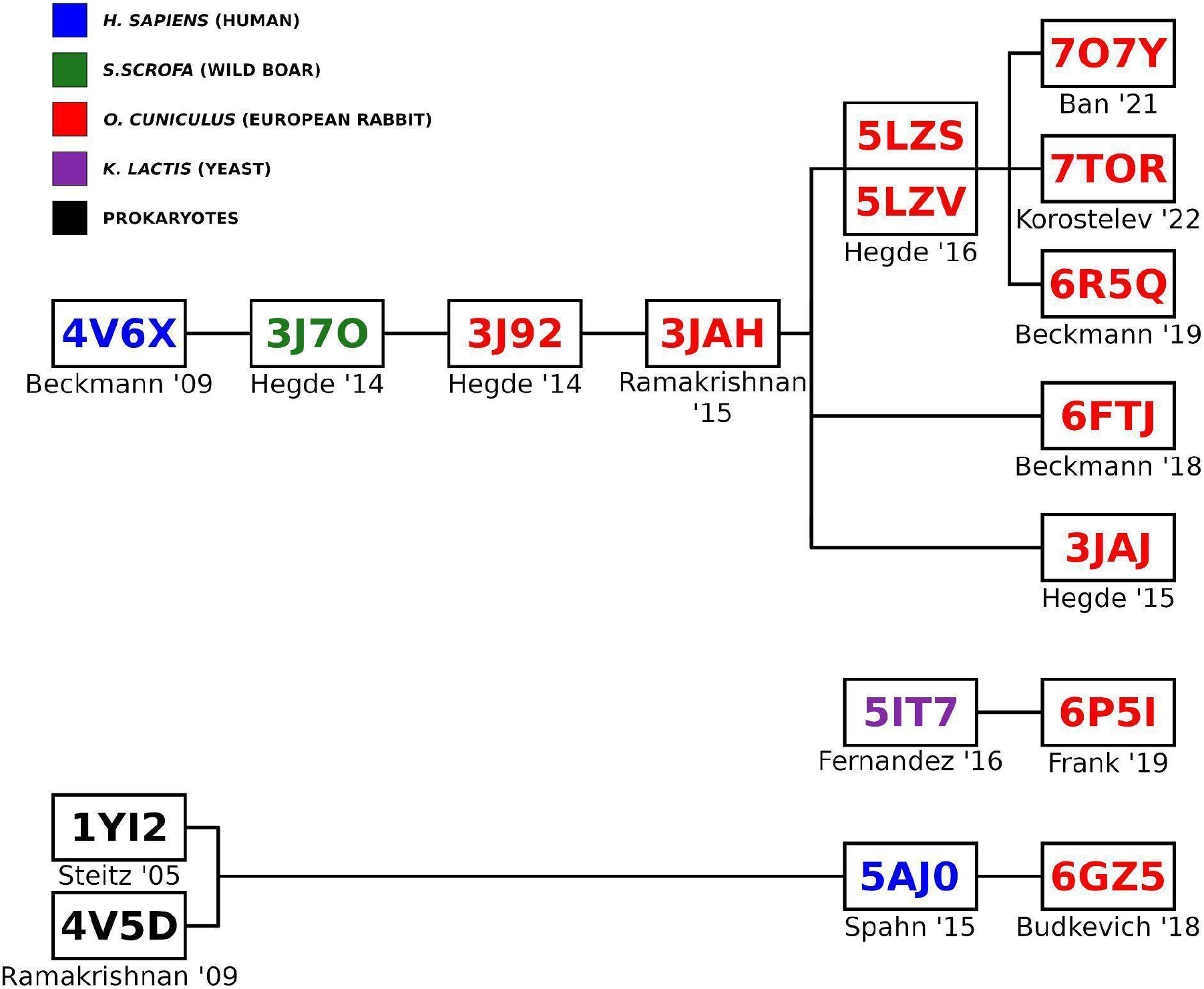
Origins of rabbit 80S ribosome Protein Data Bank entries.

Our analysis also found that some PDB entries labeled as rabbit contained sequences from other species, such as humans or boars. For example, the 28S rRNA sequence of PDB 3J92, categorized as rabbit, was derived from a boar PDB (3J7O) without any nucleotide changes (Fig. 1; Fig. 2). While boar and rabbit 28S RNA Central template sequences are 92.59% identical (Fig. 3), this observation raises important questions about the accuracy and validity of species-specific annotations in PDB entries and the conclusions drawn from such data. Additionally, a comparison between the RefSeq 28S *H. sapiens* rRNA (NR_003287) used to build the 28S rRNA of PDB entry 4V6X and RNA Central ’s rabbit template sequence (URS00009AB771_9986) revealed a surprisingly low percent identity of 84.91%. This finding highlights the significant sequence divergence that can exist even among related mammalian species, emphasizing the necessity for precise sequence verification across all organisms.

**Figure 2.**
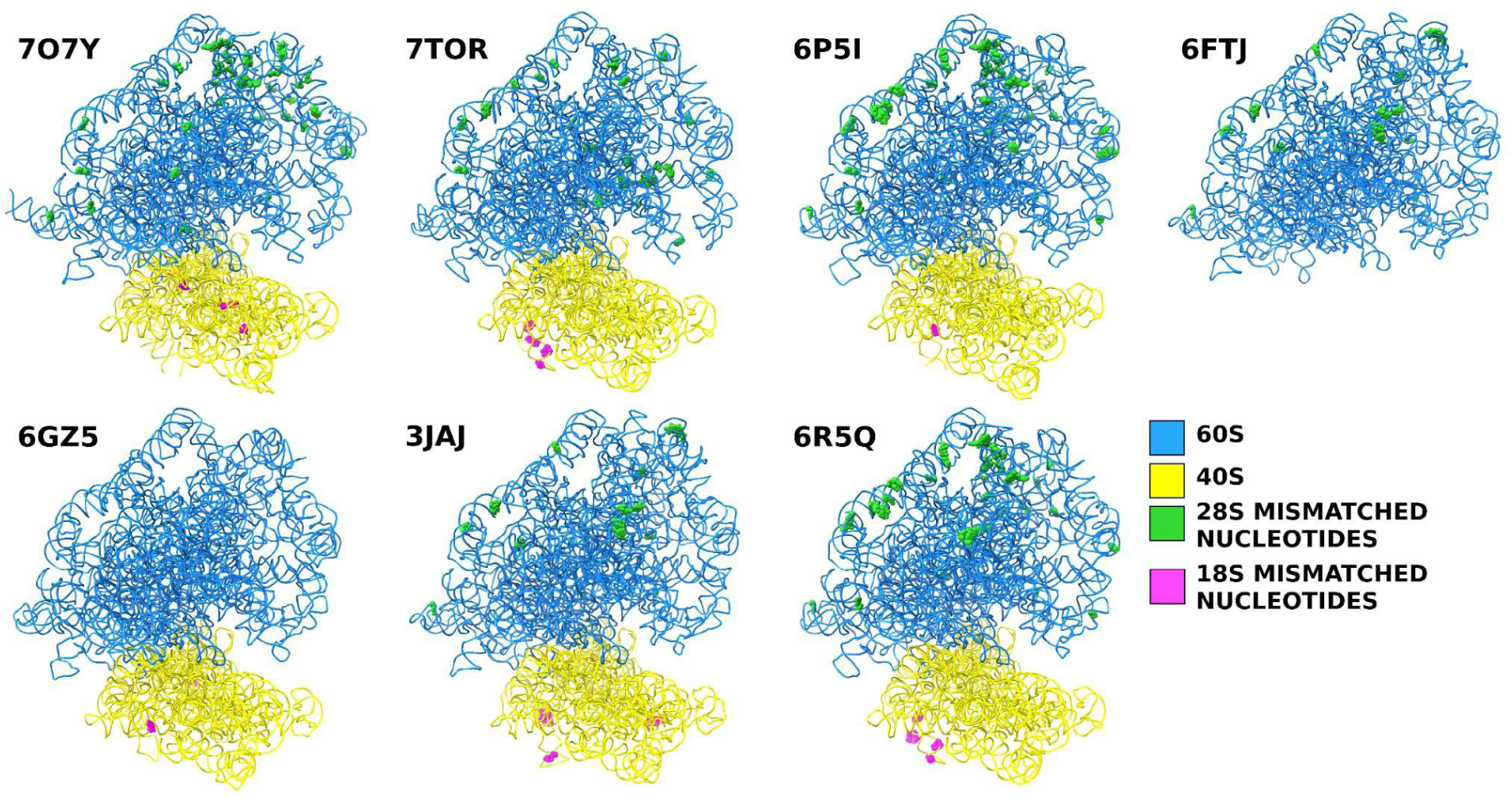
Positions of the single-nucleotide mismatches in the selected PDB entries compared to the rabbit template sequence.

**Figure 3.**
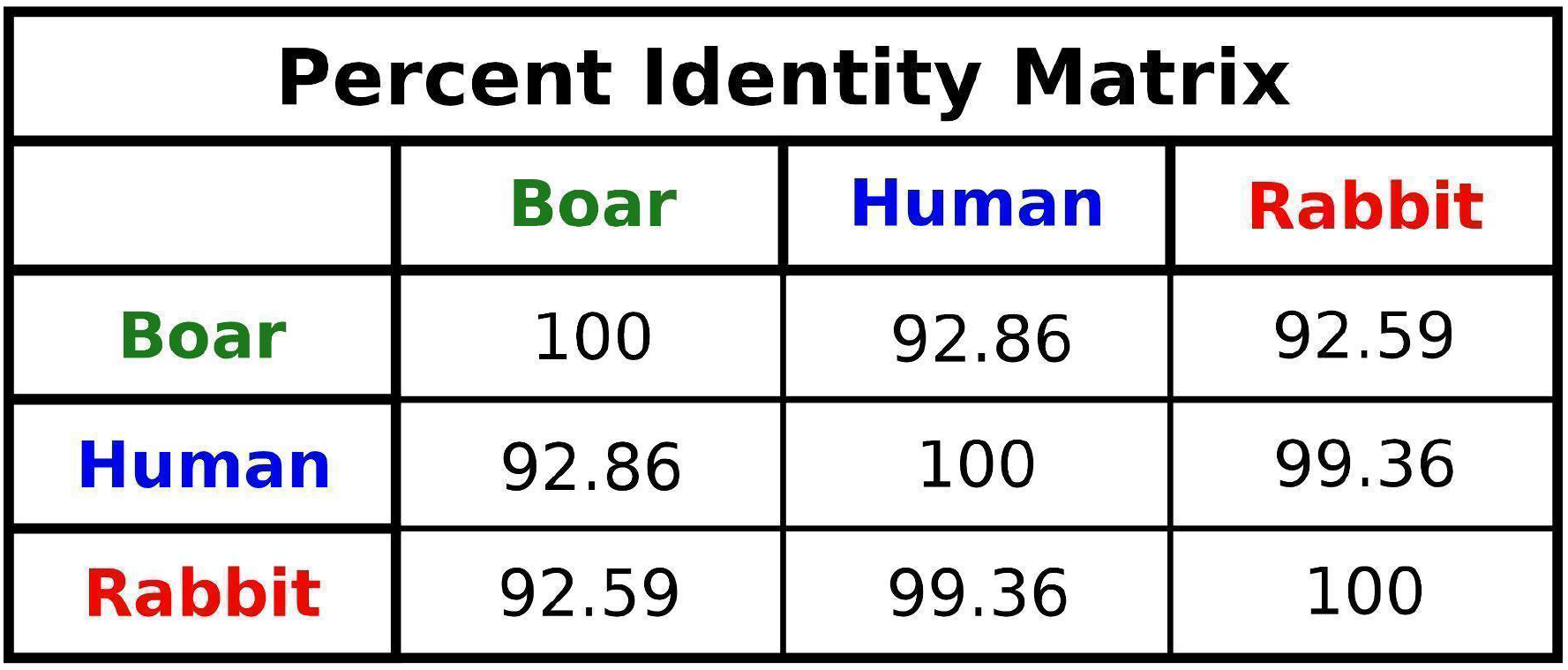
Percentage identities between the boar, human, and rabbit 28S rRNA sequences.

### Discrepancies among published 18S rRNA sequences

During all stages of ribosome function, the aminoacyl (A), peptidyl (P), and exit (E) sites on the 40S subunit are essential for the precise translation of mRNA. These sites are composed of both ribosomal protein residues and nucleotides from the 18S rRNA. Specific nucleotides of the 18S rRNA (as detailed in Table 2) are crucial for positioning tRNA, ensuring accurate codon-anticodon base pairing, interacting with initiation factors, and forming binding sites for viral IRES sequences (Pisarev et al., 2006; Abaeva et al., 2020; Simonetti et al., 2016; Quade et al., 2015; Yamamoto et al., 2015). For instance, the Kozak interaction highlights the significance of the +4 nucleotide interacting with A1819 (or A1825) of the 18S rRNA (Pisarev et al., 2006; Simonetti et al., 2020). The positioning of tRNA involves C1331 for the A-site tRNA and C1701/U1248 for the P-site tRNA in humans (Anger et al., 2013), with interactions of DHX29 with helices h16, h17, and h34 of 18S rRNA, the Sec-insertion domain of SBP2 binding to h33 of 18S rRNA, tRNA ASL with G1575 and A1576, mRNA with G1150, and additional interactions described by Petrychenko et al. for initiator tRNA with nt 1248 and 1701, and mRNA with nt 626, 961, 1207, and 1701 (Hussain et al., 2014; Simonetti et al., 2020; Sweeney et al., 2021; Hilal et al., 2022; Petrychenko et al., 2024) and engage with sense or stop codons (Brown et al., 2015; Matheisl et al., 2015). Furthermore, 18S rRNA nucleotides are crucial for binding and stabilizing IRES structures, such as U1115 and C1116 of ES7 in 18S rRNA interacting with A136 and G266 of the HCV IRES (Yamamoto et al., 2015; Quade et al., 2015; Brown et al., 2022), while C1701 and U1830 of 18S rRNA bind A1058 and U1248 of the HalV IRES (Abaeva et al., 2020). Additional interactions involving A1824-A1825/G626 of 18S rRNA and A3760/A4255 of 28S rRNA with the Israeli Acute Paralysis Virus IRES (Acosta-Reyes et al., 2019) have also been reported.

**Table 2.**
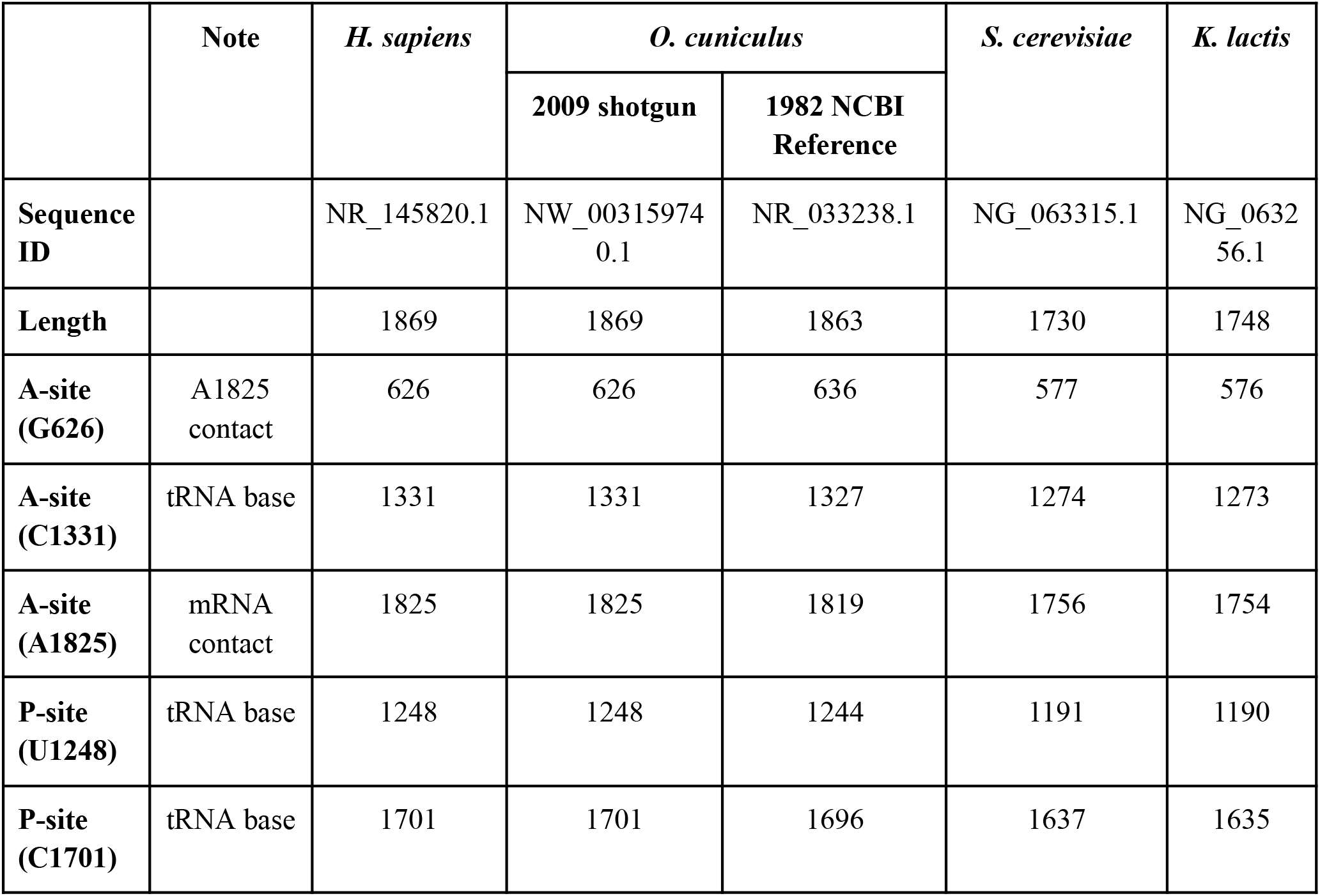
Features of the ribosomal 18S rRNA for the most commonly available models in the Protein Data Bank.

|  | Note | *H. sapiens* | *O. cuniculus* |  | *S. cerevisiae* | *K. lactis* |
| --- | --- | --- | --- | --- | --- | --- |
|  |  |  | 2009 shotgun | 1982 NCBI Reference |  |  |
| **Sequence ID** |  | NR_145820.1 | NW_00315974<br>0.1 | NR_033238.1 | NG_063315.1 | NG_0632<br>56.1 |
| **Length** |  | 1869 | 1869 | 1863 | 1730 | 1748 |
| **A-site (G626)** | A1825 contact | 626 | 626 | 636 | 577 | 576 |
| **A-site (C1331)** | tRNA base | 1331 | 1331 | 1327 | 1274 | 1273 |
| **A-site (A1825)** | mRNA contact | 1825 | 1825 | 1819 | 1756 | 1754 |
| **P-site (U1248)** | tRNA base | 1248 | 1248 | 1244 | 1191 | 1190 |
| **P-site (C1701)** | tRNA base | 1701 | 1701 | 1696 | 1637 | 1635 |

In terms of the sequence length in modeled human or yeast ribosomes, the 18S rRNA appears to be consistent, while in rabbit ribosomes, the 18S rRNA length is variable, either matching the human 18S rRNA (Genbank accession no. NR_145820.1; 1869 nucleotides; 45 models), the 1982 NCBI Reference Sequence for rabbit 18S rRNA (Genbank accession no. NR_033238.1; 1863 nucleotides; 7 models), or—in the case of the current highest-resolution rabbit ribosomal structure available—the human sequence, though out of register by +1 (1870 nucleotides; 4 models). While slightly different sequences do not impact the biological interpretation of these ribosome complexes, they do introduce ambiguity in analysis across papers as identical features are referred to using different identifiers (see Table 2).

To determine the correct 18S rRNA sequence, we examined all publicly available rabbit ribosome structures determined using cryo-EM at better than 4.0 Å resolution. While doing so, we noticed that the 1982 NCBI Reference Sequence for the rabbit 18S rRNA is partially incorrect (see supplementary Table 1 for the complete list). All cryo-EM density maps reveal at least two nucleotides missing from the 1982 NCBI Reference Sequence that are present in the human sequence (G183 and C1513). Although the nucleotide sequence of helix 18 (h18) is not entirely conserved across all species, it contains seven sequence clusters, sometimes referred to as h18 universals (Noller et al., 2022). Helix 18 has an inversion of bases in the 1982 NCBI rabbit 18S rRNA sequence. The bases G613 and C614 are reportedly reversed for the rabbit sequence (Connaughton et al., 1984; Rairkar et al., 1988), but examination of high-resolution ribosome maps (e.g., Bhatt et al., 2021) shows apparent density supporting the GC ordering seen in humans and other eukaryotic species. Together, this suggests that the sequence identity and length of the 1982 NCBI Reference Sequence for the rabbit 18S rRNA—which has been used for some cryo-EM models, biochemical experiments on rabbit ribosomes, and primer design—is partially incorrect (e.g., Pisarev et al., 2006; Simonetti et al., 2020).

The Broad Institute’s whole-genome shotgun sequencing of *Oryctolagus cuniculus* (OryCun2.0) reported a sequence that shares 99.5% identity with the human 18S, having only two nucleotide gaps. This shotgun sequence matches the human 18S sequence in length (1869 nucleotides), includes the conserved GC sequence in h18, and has both G183 and C1513 that are visibly present in all rabbit cryo-EM maps.

We used the Broad Institute’s *Oryctolagus cuniculus* Thorbecke inbred whole-genome shotgun sequence to analyze rabbit 18S rRNA sequences (OryCun2.0). This template sequence provides a highly accurate baseline for comparing the 18S rRNA sequences reported in various PDB structures. The 18S template sequence we used is found in RNA Central under the accession code URS00006F07B6_9986.

To systematically evaluate these discrepancies, we aligned six 18S rRNA sequences from published PDBs with the Broad Institute ’s rabbit sequence as a reference template. This comparative analysis exposed a range of inaccuracies, including single nucleotide substitutions and substantial insertions and deletions. For this report, we have concentrated on single nucleotide variations, setting aside issues related to insertions and deletions. This decision was driven by the realization that addressing these insertions and deletions would compromise structural integrity and make accurately remodeling the affected segments extremely challenging, even with the presently available structural information.

The sequences from these PDB entries are predominantly consistent with the reference template. For instance, the 18S rRNA sequence from PDB entry 3JAJ is 99.37% identical to the reference template, with eleven single nucleotide differences observed.

Similarly, PDB entry 6GZ5 shows an even higher level of sequence identity, at 99.82%, with only two single nucleotide variations from the template. In contrast, PDB entry 6P5I aligns with the template at 99.63%, revealing nine single nucleotide changes. PDB entry 6R5Q presents a sequence identity of 99.41%, with six discrepancies. Furthermore, PDB entry 7O7Y exhibits a 99.84% identity with just three nucleotide differences. Lastly, PDB entry 7TOR shows a sequence identity of 99.41%, with six single nucleotide differences from the template. While relatively minor, these variations are noteworthy and underscore the necessity for precision in structural biology studies.

Overall, the different PDB entries have very high sequence identities, ranging from 99.37% to 99.84%. Although there are only a few discrepancies, we must note that high accuracy is critical for precise structural interpretations and functional analyses, as minor differences can have significant implications for understanding the structure and function of rRNA within the ribosome.

This study emphasizes the importance of using accurate reference sequences for comparative analysis in structural biology. The minimal variations observed across these PDB entries suggest that the 40S structures are generally well-aligned with the reference template. However, careful attention to the remaining discrepancies is essential for ensuring the accuracy of ribosome models and advancing our knowledge of ribosomal mechanisms and functions.

### Discrepancies among published 5.8S and 5S sequences

We also used 5.8S (URS00006CE1FB_9986) and 5S (URS00006C8ED4_9986) template sequences from RNA Central to conduct a similar analysis. The 5.8S sequences in the published PDB entries matched the template sequence, with only a few indels at both ends. In contrast, the 5S rRNA showed several single nucleotide variations as well as indel discrepancies. Notably, the 5S template sequence consists of 144 nucleotides, while the PDB entries contain only 119 nucleotides.

## Conclusion

The discrepancies we have found have profound implications for structural biology. Inaccurate and inconsistent rRNA sequences in published PDB entries can lead to misinterpretations of ribosome structure and function, potentially affecting the validity of many downstream research findings and applications. The propagation of incorrect sequences through successive studies exacerbates these issues, leading to a cascade of errors and misannotations. To address these challenges, we propose that all future rabbit ribosome models use the numbering and sequence from the Broad Institute’s rabbit genome sequencing project. Although rabbits possess multiple cistrons for their rRNAs just like many other organisms, potentially introducing sequence variability (Martin-DeLeon, 1980; Hori et al., 2023; Rothschild et al., 2024), our study provides a crucial foundation for understanding ribosome structure and function by establishing a consensus sequence. In this way, we believe our study enables more accurate and meaningful interpretations of structural results, ultimately advancing our understanding of ribosome biology. We are also depositing the sequence-corrected 28S and 18S structures into the RCSB PDB to facilitate the use of correct sequences by the field. This will also maintain consistency in numbering important ribosomal subunit sites between rabbit and human ribosomes. By adopting rigorous sequence verification and correction practices and utilizing trusted template sequences,we can ensure more accurate and reliable models, advancing our understanding of ribosome structure.

## Methods

For this study, we selected seven PDB entries (7O7Y, 7TOR, 6P5I, 6FTJ, 6GZ5, 3JAJ, and 6R5Q) from seven different published studies. We utilized the curated RNA Central entry URS00009AB771_9986 as a consensus sequence for the rabbit 28S rRNA template, which is accessible under GenBank accession AAGW00000000.2, Rfam accession RF02543, and ENA accession GL019111 (Broad Institute’s rabbit whole genome sequencing trials (OryCun2.0)). The 28S rRNA sequences from each PDB were downloaded from RCSB and aligned against the consensus sequence using Clustal Omega (EMBL-EBI) (Sievers et al., 2011). This alignment was reviewed for single-nucleotide discrepancies, which were meticulously recorded. Insertions and deletions were excluded from this analysis. The 18S, 5S, and 5.8S rRNA sequences were also downloaded and aligned against the consensus 18S (URS00006F07B6_9986), 5S (URS00006C8ED4_9986), and 5.8S (URS00006CE1FB_9986) rRNA sequences. We used

ChimeraX (Meng et al., 2023) to examine structural details, verify alignment fidelity, and resolve any alignment issues. Additionally, Coot (Emsley and Cowtan, 2004) was used to address and correct sequence errors as they arose during analysis. Finally, each refined structure was further refined against our unpublished *Oryctolagus cuniculus* 80S ribosome maps and already published *Oryctolagus cuniculus* 40S maps (Brown et al., 2022) to ensure rabbit-specific ribosomal conformation using phenix.refine (Afonine et al., 2012). This step provided an additional level of precision to our analysis.

## Acknowledgments

This work was supported by grants from the National Institutes of Health: R35 GM139453 to J.F., R35 GM122602 to T.V.P., and R01 GM097014 to C.H. We also would like to thank the Irvington High School Science Research Program (Irvington, New York).

## Notes

### Competing Interest Statement

The authors have declared no competing interest.

### Summary of Updates

Figure 2 colors were changed for better visibility. Rest are unchanged.

